# The discriminant pattern of pleural fluid inflammatory mediators between tuberculosis and other causes of exudative pleural effusion

**DOI:** 10.1101/667360

**Authors:** Vinícius da Cunha Lisboa, Raquel da Silva Corrêa, Marcelo Ribeiro-Alves, Isabelle Ramos Lopes, Thiago Thomaz Mafort, Ana Paula Gomes dos Santos, Thaís Porto Amadeu, Rogério Lopes Rufino Alves, Luciana Silva Rodrigues

## Abstract

Pleural tuberculosis (PlTB), a form of extrapulmonary TB, remains as a challenge in the diagnosis among many causes of pleural effusion. We recently reported that the combinatorial analysis of interferon-gamma (IFN-γ), IFN-γ-inducible protein 10 (IP-10), and adenosine deaminase (ADA) from the pleural microenvironment was useful to distinguish pleural effusion caused by TB (microbiologically or not confirmed cases) among other etiologies. In this prospective cohort study, a set of inflammatory mediators was quantified in blood and pleural fluid (PF) from exudative pleural effusion cases, including PlTB (n = 22) and non-PlTB (NTB; n= 17) patients. The levels of IL-2, IL-4, IL-6, IL-10, IL-17A, IFN-γ, TNF, IP-10, TGF-β1, and ADA were measured and a principal component analysis was applied in order to identify the mediators who contributed most for the variance in data. IFN-γ, IP-10, TNF, TGF-β, and ADA quantified in PF showed significantly higher concentrations in PlTB patients when compared to NTB ones. When blood and PF were compared, we have identified significantly higher concentrations of IL-6 and IL-10 in PF, in both groups. TGF-β, solely, showed significantly increased levels in PF and blood from PlTB when both clinical specimens were compared to NTB patients. Principal components analysis from PF revealed that the ADA, IP-10, TGF-β, and IFN-γ contributed most for the discriminatory capacity between TPlB and NTB. Our findings showed that important inflammatory mediators in PF may discriminate TB cases from other causes of exudative effusion, the main diseases considered in the differential diagnosis of PlTB.

## INTRODUCTION

Tuberculosis, caused by *Mycobacterium tuberculosis* (Mtb), is currently endemic in the world and represents an important public health problem every year. Globally, in 2017, more than 10 million new cases of TB were reported with an estimated 1.3 million deaths. Among infectious diseases, TB is the leading cause of death from a single agent, surpassing the human immunodeficiency virus (HIV) infection (1). Although TB affects mainly the lungs, extrapulmonary forms can appear as an initial manifestation in approximately 25% of adults with TB, of which the pleural space is the second site of involvement followed only by the lymph nodes (2). In Brazil, a high burden TB country, PlTB is responsible for more than 40% of cases among many clinical sites of extrapulmonary TB (3) and still imposes a challenging diagnosis due to, mainly, its paucibacillary nature and the need of invasive producers (4).

Cellular immune response (Th1 immunity) involving CD4+ T-lymphocytes, classically studied and associated with the containment of Mtb in pulmonary parenchymal TB, is also predominant in TB pleuritis, which is confirmed by the higher levels of interferon-gamma (IFN-γ) and other inflammatory cytokines (e.g., IL-12) in pleural fluid in comparison to peripheral blood (2, 5–7). IFN-γ promotes cell differentiation, stimulates an increased phagocytic activity and intermediate nitrogen and oxygen species production, which are bactericidal and participate in resistance to Mtb infection (8, 9). In addition, other T-cell effectors patterns are involved in Mtb control in the pleural microenvironment, such as Th17, which express the retinoic acid-related orphan receptor gamma t (RORγt), and are characterized by secretion of large quantities of IL-17 (also known as IL-17A), IL-21, and IL-22 (9, 10). Th17 cells induce the expression of many pro-inflammatory factors, chemokines, ultimately involved in granulopoiesis and recruitment of innate cells, mainly neutrophils, especially in the early stages of infection (11, 12).It is well described that patients at early stages of the PlTB (less than 2 weeks duration) or those who present pleural effusion with high complexity (e.g., loculated pleural effusion, TB empyema) are more likely to have a neutrophilic exudate (reviewed by 13), which may contribute to injuries and decreases pleuro-pulmonary functions.

Since that the gold standard for the diagnosis of PlTB which is the detection of Mtb in the sputum, pleural fluid or pleural biopsy has a discrete and variable yield, the histological demonstration of caseating granuloma even in the absence of acid-fast bacilli can be sufficient for anti-TB treatment (14, 15). Additionally, values > 40 IU/L of adenosine deaminase (ADA) in pleural effusion, a purine-degrading enzyme, associated with a predominantly lymphocytic exudate, and clinical suspicious of TB, altogether, indicates that the most likely diagnosis is tuberculosis (16, 17). However, high pleural fluid ADA values can also be found in certain conditions, such as adenocarcinoma, lymphoma, rheumatoid arthritis, and pleural empyema of bacterial etiology, making the differential diagnosis very hard (18, 19).

Considering the difficulty in the differential diagnosis already mentioned and the current knowledge about the products of the immune response against Mtb in pleural space, the present study aimed to identify biomarkers among Th1, Th2, and Th17 T-cells subsets and other inflammatory mediators in peripheral blood and pleural fluid which could present high potential of utility for the PlTB diagnosis among exudative pleural effusion from other etiologies. Based on a principal component analysis (PCA), we could demonstrate a pattern of mediators which was able to discriminate TB from non-TB pleural effusion.

## MATERIAL AND METHODS

### Study population and settings

Patients aged ≥ 18 years with pleural effusion under investigation with thoracentesis indication were recruited in this cross-sectional prospective study which was conducted at the Pulmonology and Tisiology Service, Pedro Ernesto University Hospital/Rio de Janeiro State University (HUPE/UERJ), a tertiary care center at Rio de Janeiro, RJ, Brazil. Patients who were under 18 years of age, pregnant, or refused consent were not recruited. Of 49 recruited patients, 10 were excluded: 8 patients had transudative pleural effusion (cardiac or renal failure), and 2 patients were HIV-seropositive. Thus, 39 patients with exudative pleural effusion were enrolled in the study: 22 PlTB and 17 non-TB (NTB) patients. ***PlTB cases*** were defined by the patient history reviewed, followed by a detailed physical examination, and at least one diagnostic criteria: i) positive results in the microbiological and/or histopathological tests (acid-fast bacilli smear microscopy, mycobacterial culture, or Xpert MTB/RIF^®^) on pleural fluid or pleural tissue; ii) presence of granuloma with or without caseous necrosis; iii) clinical manifestations suggesting TB (fever, pain, dyspnea, cough, night sweats, hyporexia, and/or weight loss) in combination with a lymphocytic pleural effusion, followed by a full recovery after at least six months of anti-TB treatment. ***Non-TB cases*** consisted of patients with pleural or pleuro-pulmonary diseases, excluding active TB based on clinical, laboratory, radiological, microbiological and/or pathological features. Malignant pleural effusions were diagnosed by a positive pleural fluid cytology result or malignant cells identified in the pleural fragment. Even when both of these tests results were negative, malignant effusion was diagnosed when a primary cancer was known to have disseminated and no other cause of pleural effusion was identified. Patients who did not fit the criteria used for PlTB diagnosis as above and with unknown cause of pleural effusion were classified as “undefined” pleural effusion and considered as non-PlTB. Medical information, peripheral blood, and pleural fluid sample collection were obtained from all study subjects after signing a written consent. The study protocol was approved by the respective institutional ethics committee (HUPE/UERJ; number 1.100.772).

### Sample collection

Ultrasound-guided thoracentesis was performed by a trained pulmonologist who collected pleural fluid which was directly drawn into collection tubes for routine diagnostic tests, including chemistry panel, total, and differential cell count, ADA measurement by Hermes Pardini laboratory according Giusti’s method (20), cytopathology, microbiological analysis (bacteria, fungi and mycobacteria), and inflammatory biomarkers for the purpose of the present study. During collection, whole blood and pleural fluid were sampled in appropriated collection tubes without anticoagulant additive. After collection, whole blood and pleural fluid tubes were centrifuged at 1000 x g for 10 min and 25 °C or 4 °C, respectively. Then, serum and pleural fluid (without cells) samples were aliquoted and stored frozen at −20 °C until cytokines quantification.

### Cytokines assays

Cytokine levels in clinical samples were assessed using the following commercially available kits: i) human Th1/Th2/Th17 Cytokine Kit (BD Bioscience, San Jose, CA, USA) based on the principle of cytometric bead array (CBA) technology for simultaneous detection of seven cytokines (IL-2, IL-4, IL-6, IL-10, TNF, IFN-γ, and IL-17A). Briefly, capture beads labeled with distinct fluorescence intensity (allophycocyanin; APC) conjugated to specific antibodies for cytokines were incubated around 3 hours in the dark at room temperature with the undiluted samples, and fluorescent detection antibody (phycoerythrin; PE). All unbound antibodies were washed and samples acquired on a BD fluorescence-activated cell sorting (FACS) analyzer FACSCanto II. Cytokine standard curves ranged 0-5,000 pg/mL. ii) IP-10 and TGF-β levels were measured by enzyme-linked immunosorbent assay (ELISA) sandwich using human CXCL-10/IP-10 DuoSet ELISA (R&D Systems Inc, MN, USA) and human/mouse TGF beta 1 ELISA Ready-SET-Go! Kit (2^nd^ Generation; Affymetrix, eBioscience), respectively, following the manufacturer’s instruction. The range of these assays was 31.3-20,000 pg/mL for IP-10 and 15.6-1,000 pg/mL for TGF-β. Readings greater than the upper limit were set at 20,000 (IP-10) or 1,000 (TGF-β) pg/mL for the purpose of analysis.

### Statistical analysis

For the description of the population included in the study, according to their sociodemographic and clinical characteristics among the individuals with exudative pleural effusion due to PlTB or other causes (non-TB), non-parametric Mann-Whitney test were used for continuous variables or Fisher’s exact tests for comparison of the relative frequencies of the different levels of nominal/categorical variables. In the comparison between the levels of log-transformed expression (bases 10) of proteins in peripheral blood/serum and pleural fluid (tissue effect) between individuals with or without TBPl (TB effect), the expected mean marginal values obtained from multiple linear regression (log-linear) models of fixed effects were used with the inclusion of first-order interactions between the main tissue and TB effects. For the adjusted models, graphical analysis of residuals was performed to confirm their randomness. In the comparisons between expected mean marginal values obtained from linear regression models, adjustments of the confidence level were made by Sidak’s method, and p-value adjustments by multiple comparisons by Tukey’s method. Finally, for log10-transformed protein and ADA expression data, a multivariate principal component analysis (PCA) was performed to visualize the distribution of sample individuals in 2D dimensional spaces. Ellipses of the quantiles 68% of the normal distribution adjusted to the individuals of the different interest groups in these new dimensional spaces are presented. The level of significance, P ≤ 0.05, was used in the analysis, and all analyses were performed in R software version 3.5.2.

## RESULTS

### Patients and characteristics

Study population was composed by 39 individuals who were diagnosed as PlTB (n = 22) or non-TB (n = 17) according previously described. Their sociodemographic and clinical data are shown in Table 1. We observed a significant difference between the age distributions between the groups, which presented medians corresponding to 65 years (IQR: 20) in the non-TB group, and 41 years (IQR: 14) in the PlTB group (p < 0.0001). Smoking and alcoholic habits among participants did not show statistical differences. Similarly, symptoms presentation was not dissimilar among groups. Fourteen (82%) patients in the non-TB group had one or more comorbidities, showing that this group had a significantly higher number of patients with comorbidities than observed in the P1TB group, which had 4 individuals (18%) with others diseases (p = 0.0217). The most prevalent comorbidity was hypertension, which was reported in 6 (35%) non-TB patients and 2 (9%) P1TB patients. Among non-TB patients, 13 were malignancies, 1 autoimmune disease (systemic lupus erythematosus), and 3 undefined pleural effusion. PlTB, Pleural tuberculosis; PE, pleural effusion; IQR, Interquartile range. Values expressed as n (%; from the total population) unless otherwise stated.

**Table 1.** Sociodemographic and clinical characteristics of the study population.

| Characteristics /Group | Non-TB<br>(n=17) | PITB<br>(n=22) | p-value |
| --- | --- | --- | --- |
| Age, years median (IQR) | 65 (14) | 41 (20) | 0.0001 |
| **Gender (%)** |  |  |  |
| Female | 7 (17.9) | 8 (20.5) | 1 |
| Male | 10 (25.6) | 14 (35.9) |  |
| **Current smoker (%)** | 2 (5.1) | 3 (7.7) | 0.2836 |
| **Alcohol use (%)** | 2 (5.1) | 9 (23.1) | 0.1032 |
| **Comorbidities (%)** | 10 (25.6) | 4 (10.3) | 0.0172 |
| Hypertension | 6 (15.4) | 2 (5.1) | 0.059 |
| Diabetes | 3 (7.7) | 0 (0) | 0.0744 |
| Cardiac Insufficiency | 3 (7.7) | 0 (0) | 0.0744 |
| Hepatitis | 2 (5.1) | 1 (2.6) | 0.5703 |
| **Symptoms (%)** |  |  |  |
| Fever | 3 (7.7) | 9 (23.1) | 0.1612 |
| Cough | 13 (33.3) | 9 (23.1) | 0.0516 |
| Chest Pain | 5 (12.8) | 10 (25.6) | 0.5 |
| Dyspnea | 13 (33.3) | 12 (30.8) | 0.3068 |
| Night Sweats | 3 (7.7) | 4 (10.3) | 1 |
| **Definitive cause of PE (%)** |  |  |  |
| Tuberculosis |  | 22 (56.4) |  |
| Malignancy | 13 (33) |  |  |
| Autoimmune disease | 1 (2.5) |  |  |
| Undefined | 3 (7) |  |  |

PITB, Pleural tuberculosis; PE, pleural effusion; IQR, Interquartile range. Values expressed as n (%; from the total population) unless otherwise stated.

### Cytokines measurement in blood and pleural fluid from PlTB and non-TB patients

In order to evaluate the potential diagnosis of cytokines Th1/Th2/Th17, IP-10 chemokine, and ADA in exudative cases of pleural effusion, serum and pleural fluid samples from PlTB and non-TB patients were analyzed. As recently reported by our group (7) and others (21–23), IFN-γ and IP-10 levels were significantly increased (p < 0.0001 in both) in pleural fluid comparison to serum in PlTB group (Figure 1A and H). As shown in Figure 1B, TNF concentration also showed a significant increase in the pleural fluid when compared to serum in PlTB patient (p = 0.0016).

**Figure 1.**
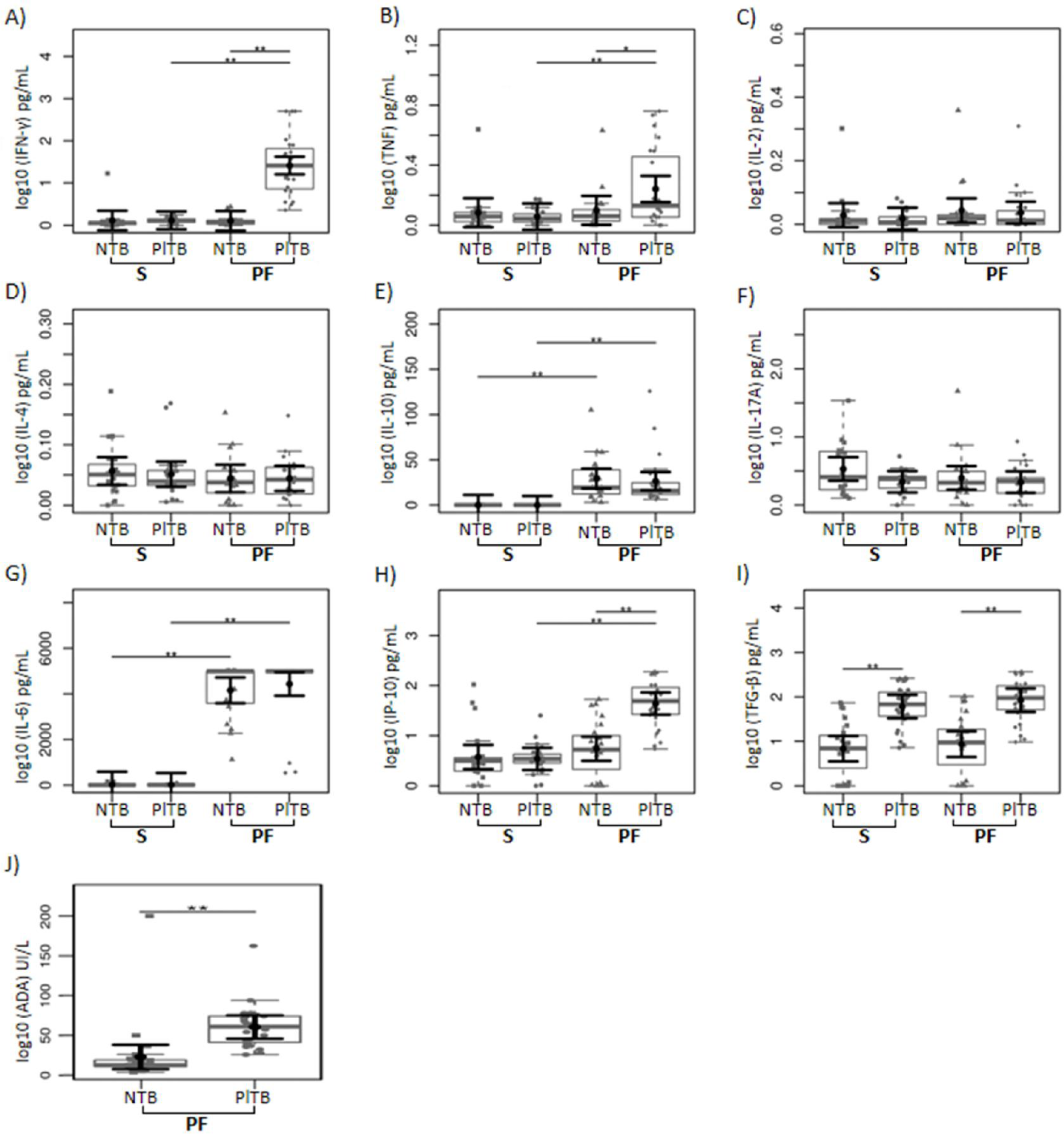
Cytokines and ADA levels in serum and pleural fluid from PlTB and NTB patients. Cytokines were dosed by CBA (IL-2, IL-4, IL-6, IL-10, TNF, IFN-γ, and IL-17A). The levels obtained from each cytokine were analyzed on a logarithmic (base = 10) scale and illustrated using boxplots to compare serum (S) and pleural fluid (LP) data between the non-TB (NTB) and TBPl groups. The small grey dots represent individual cases and the boxplots represent the interquartile range and the median of the sample (solid grey central line). Larger black dots and vertical bars represent expected mean marginal values estimated by the linear model and its 95% confidence intervals (95% CI). Comparisons of means between groups were performed by contrasts/differences obtained after linear bi and multivariate models, adjusted by regressions by ordinary least squares. * p < 0.05; ** p < 0.01.

When these cytokines were compared with discriminatory objectives between PlTB and non-TB patients, we predominantly observed significant differences in pleural fluid. IL-6 and IL-10 levels presented the same behavior when serum and pleural fluid were compared in PlTB or non-TB groups (Figure 1G and E, respectively). Both IL-10 and IL-6 concentrations show that patients in both PlTB (p < 0.0001 in both) and NTB (p < 0.0001 in both) groups show increased concentrations of this cytokine in pleural fluid when compared to serum in their respective groups. As expected, ADA levels were significantly higher in pleural fluid of PlTB patients compared to non-TB (p < 0.0001). Interestingly, TGF-β concentrations were significantly higher in the serum (p < 0.0001) and pleural fluid of P1TB patients, compared to concentrations found in non-TB patient samples (p < 0.0001). Concentrations of this growth factor showed no significant serum and pleural fluid difference when compared in the same group (Figure 1I).

Finally, IFN-γ, TNF, IP-10, TGF-β and ADA concentrations in the pleural fluid presented a differentiated profile between PlTB and non-TB patients. Cytokines IL-17A, IL-4, and IL-2 did not show significant differences in their concentrations.

### Principal component analysis of pleural fluid cytokines

Finally, it was observed whether the overall cytokines profile was able to discriminate PlTB and non-TB cases. In the principal components analysis (PCA), 57.21% of the total variance in response to 9 cytokines and biomarkers was expressed by 2 principal components (Table 2; Figure 2). The first component accounted for a total of 40.58%, while the second accounted for 16,63% of the total variance (Figure 2). Altogether, these 9 cytokines were able to discriminate between PlTB and non-TB. The most determinant variables of each of these two principal components were respectively ADA, IP-10, TGF-β, IFN-γ, and TNF, for the first principal component (PC1), and IL17A, IL-4, and IL-2 for the second principal component (PC2).Principal component analysis of inflammatory biomarkers in pleural fluid from patients with pleural effusion by pleural tuberculosis and other diagnoses. Shaded values represent the most important biomarkers in the component definition. NA: not applicable.

**Table 2.** Principal components analysis.

| Component | Eigenvalue | Difference | Proportion | Cumulative |
| --- | --- | --- | --- | --- |
| Component 1 | 3.95 | 2.33 | 40.58 | 40.58 |
| Component 2 | 1.62 | 0.31 | 16.63 | 57.21 |
| Component 3 | 1.32 | 0.42 | 13.50 | 70.70 |
| Component 4 | 0.90 | 0.10 | 9.22 | 79.92 |
| Component 5 | 0.79 | 0.30 | 8.16 | 88.08 |
| Component 6 | 0.50 | 0.15 | 5.08 | 93.16 |
| Component 7 | 0.35 | 0.10 | 3.55 | 96.71 |
| Component 8 | 0.24 | 0.16 | 2.48 | 99.19 |
| Component 9 | 0.08 | 0.07 | 0.79 | 99.98 |
| Component 10 | 0.00 | NA | 0.02 | 100.00 |

| Variable | Component 1 | Component 2 |
| --- | --- | --- |
| ADA | -0.41429 | -0.21968 |
| IP-10 | -0.44817 | -0.19145 |
| TGF-β | -0.4445 | -0.18435 |
| IL17A | -0.0793 | 0.394041 |
| IFN-γ | -0.44469 | -0.18198 |
| TNF | -0.38718 | 0.253531 |
| IL-10 | -0.1654 | 0.330127 |
| IL-6 | -0.03609 | -0.02504 |
| IL-4 | -0.16608 | 0.56734 |
| IL-2 | -0.1405 | 0.443796 |

Principal component analysis of inflammatory biomarkers in pleural fluid from patients with pleural effusion by pleural tuberculosis and other diagnoses. Shaded values represent the most important biomarkers in the component definition. NA: not applicable.

**Figure 2.**
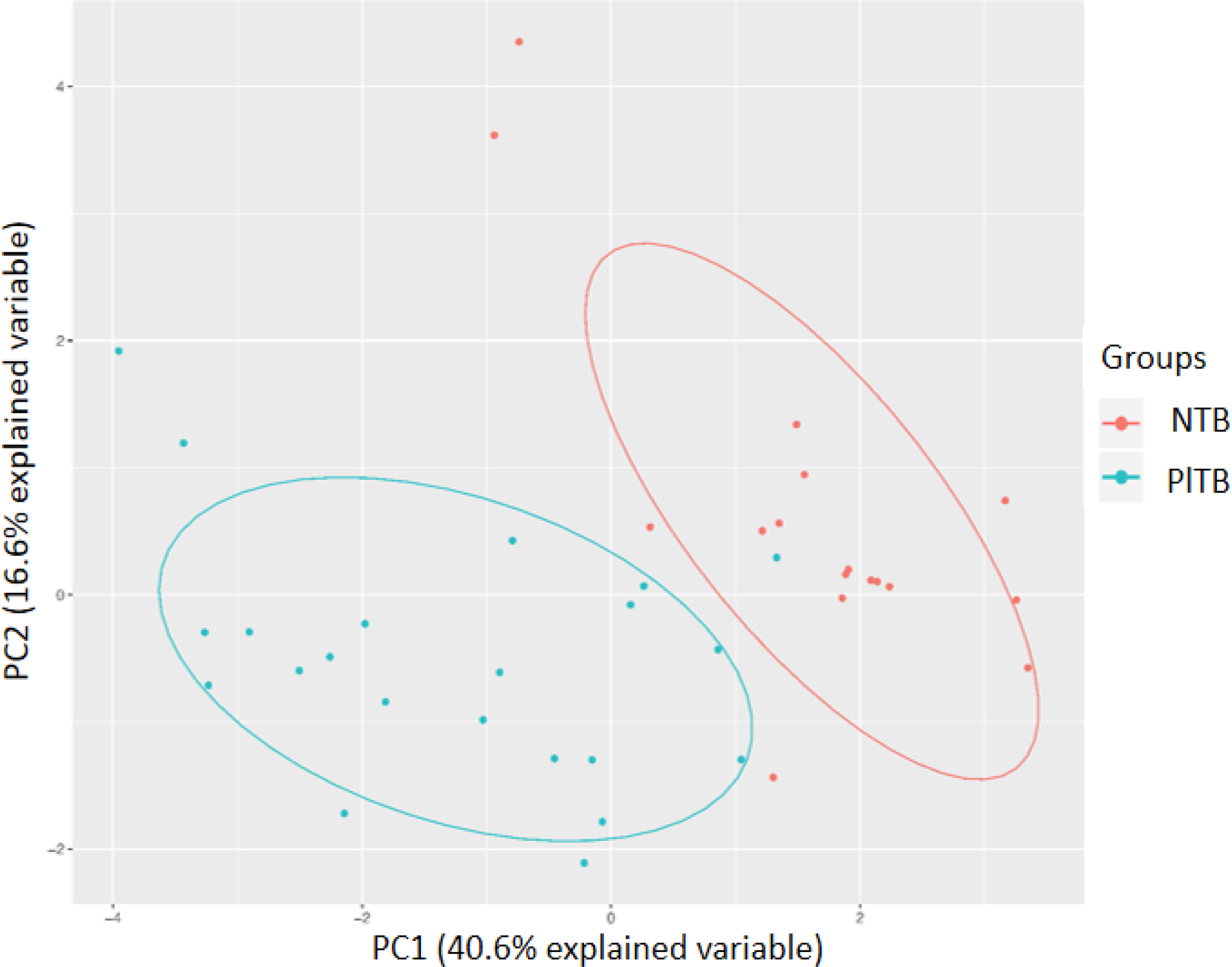
A pattern of inflammatory biomarkers in pleural fluid discriminates PlTB from NTB patients. The analysis of variance of cytokine concentrations by CBA (IL-2, IL-4, IL-6, IL-10, TNF, IFN-γ, and IL-17A) and ELISA (TGF-β and IP-10) ADA were evaluated by the PCA method, with the objective to finding heterogeneity between the cytokine profile of patients with pleural tuberculosis (P1TB) and other diagnoses non-TB (NTB). Two components were used to explain most of the total variation of these data.

## DISCUSSION

Among many known causes of pleural effusion, heart failure, malignant conditions, pneumonia, and PlTB are responsible for three-quarters of all cases (24). Currently, there is scarce literature comparing PlTB with other causes of exudative pleural effusions, which contributes to the difficulty of establishing criteria for the differential diagnosis of PlTB. The present study sought to find elements that are capable of differentiating the tuberculous effusion from other agents causing pleural effusion. Analyzing the pleural microenvironment, the quantification of cytokines showed higher concentrations of IFN-γ, TNF, TGF-β, IP-10 and ADA in LP of P1TB patients, when compared to LP of non-TB patients. Analysis of principal components revealed that these cytokines and inflammatory mediators showed the largest variations associated with a partial distinction between PlTB and non-TB patients.

As mentioned, ADA dosage is routinely used as a marker of PlTB, although it does not define the differential diagnosis (17, 18). As for the general characteristics of pleural effusions, as expected, the median value corresponding to ADA concentrations was significantly higher in the PlTB group, compared to the non-TB group. Although very useful in the differentiation of tuberculous effusion, several authors diverge about the true diagnostic value of the ADA, often setting other cutoff values. However, although they are found in greater amounts in the pleural fluid, these mediator have not been used, alone, as differential markers in exudative pleural effusions. In a recent work published by our group (7), we proposed a model where the values of IP-10, IFN-γ, and ADA in a revised cutoff value, analyzed together, can be used in the differential diagnosis of P1TB with high performance in microbiologically unconfirmed cases of PlTB.

The immune system plays a pivotal role in the evolution of Mtb infection. To contain the infection, the defense cells together with inflammatory mediators generated at the site of the infection produce a potent and aggressive response, which can generate important tissue lesions (25). In addition, the direct action on the mesothelial cells and vascular endothelium, present great participation of the tissue repair and fibrosis processes, causing functional impairment of the pleural and lungs (26, 27). At the same time, an insufficient immune response may allow the multiplication and dissemination of the bacillus (25). Otherwise, the immune system products can be used to identify pleural effusions caused by TB.

Classically, the Th1 response is the most studied in TB, being responsible both for the containment of Mtb and for the tissue injury caused by the excessive response to the bacillus (reviewed by 28). As expected, Th1-related cytokines IFN-γ and TNF, as well as the biomarker IP-10, are increased in the pleural fluid of P1TB patients compared to blood (Figure 1). Recently, the dosage of IFN-γ in pleural effusion raised importance as an auxiliary method for the diagnosis of PlTB, becoming an example of a test used for this purpose, since this cytokine is at high levels during the active phase of the disease (23, 28). The IFN-γ-release assay (IGRA) has also been highlighted in this context. The test evaluates the activity of T lymphocytes under stimulation of Mtb ESAT-6 and CFP-10 antigens. However, as reviewed by Aggarwal and collaborators (2015) there are many conflicting results regarding this diagnostic method of active tuberculosis, both in pulmonary and pleural forms (29). Moreover, as recently delineated by our group, IGRA has a poor meaning in PlTB (7), perhaps due to their paucibacillary nature or due to the enrichment of inflammatory mediators in pleural space, without needing of an additional antigen-stimuli. TNF is another important mediator in the response against Mtb and it is directly related to the maintenance of the granuloma structure, maintaining the colonization of the bacillus and necrosis area in a restricted manner (30). Other evidence shows that patients treated with an anti-TNF antibody developed active tuberculosis after reactivation of latent infection (31). In addition, TNF is important in the intracellular control of Mtb (Review by 25). Li et al. (2014) found a higher diagnostic value in TNF measurements than that found in ADA values (22). IP-10 is well studied as a possible biomarker in TB and is directly associated with INF-γ since its production is mainly induced by this cytokine. As revised by Porcel (32), IP-10 is not an essential biomarker for the PlTB diagnosis but has been the subject of several studies in this context, based on its participation in the immunopathogenesis of the disease and their correlation with IFN-γ (7, 21).

Recently, the Th17 response also gained prominence in the immunopathology of tuberculosis, especially in the early stages of infection (9, 10). Particularly in PlTB, Ye et al. (2011) showed an increase of lymphocytes with Th17 profile in pleural fluid compared to blood. Our results, presented here, show high concentrations of IL-6 and TGF-β in pleural fluid of PlTB patients compared to serum (34). These two biomarkers are critical in the differentiation of Th17 cells (35). Therefore, although our study did not focus on the characterization of Th17 cells, it is quite probable that the microenvironment, through the high concentration of IL-6, TGF-β, and the low concentrations of IL-2 is favoring the differentiation of this T-lymphocytes effector phenotype in the PLTB group.

IL-10 is a cytokine involved in the suppression of the immune response (36). In the case of TB, it has been associated with suppression of dendritic cell activity, the formation of foamy macrophages and defective formation of granuloma (37–39). The production of IL-10 is one of the more classic mechanisms of suppression by T regulatory cells, however, this cytokine can be produced by many other cells of the immune system, such as macrophages, B lymphocytes and Th2 lymphocytes (40). Our study has shown higher concentrations of IL-10 in the pleural fluid of patients with PlTB compared to serum, and in the same way in the NTB group. However, the methodology used in this study was not able to identify which cells present in the pleural fluid were responsible for the increase of IL-10 concentrations, as well as the other cytokines. Geffner et al. (2013) showed an increased IL-10 production after stimulation of mononuclear cells in pleural fluid and peripheral blood with Mtb antigens, and decrease of this cytokine after removal of culture Treg cells provides evidence that Treg is also responsible for the production of IL-10 from the pleural cavity (41).

The cytokine pattern related to the Th2 effector phenotype was also evaluated. In the methodology employed, but did not detect significant levels of IL-4. This finding confirms the literature data that show little influence of this effector phenotype in cases of tuberculosis (2, 33), although IL-4 concentrations in miliary TB have already been reported (5).

Another important finding in our study was the quantification of TGF-β in serum and pleural fluid. This growth factor, secreted by monocytes, is a chemotactic agent for fibroblasts, plays an important role in extracellular matrix remodeling (42). One of the possible contributions of TGF-β to the pathophysiology of PlTB is its ability to induce fibrosis, as shown in the study by Sasse et al (2003), where animals infected with Mtb showed increased pleural thickening in proportion to the increase in TGF-β (43). Seiscento et al (2007) also found elevated TGF-β levels in serum and pleural fluid of PlTB patients, associating with the degree of pleural thickening in these patients (44). Our findings, together with the evidence found in the literature, reinforce the hypothesis that this mediator may be related to the development of pleural effusions in TBP1 patients since TGF-β levels were found to be significantly higher in the pleural fluid of these patients, compared to the results found in non-TB patients. Although the cited studies found a significant increase of TGF-β in pleural fluid and serum, the comparison group in the experimental model of these studies was composed of patients with transudative pleural effusion. Our work was able to detect the increase of TGF-β in the serum and pleural fluid of PlTB patients, compared to blood and pleural fluid in patients with other causes of exudative effusion.This finding may contribute to future investigations, associating TGF-β as a possible biomarker to aid in the differential diagnosis of PlTB.

In fact, the cytokines analyzed alone are not able to provide data of high specificity and sensitivity, especially in comparison to exudative effusions. However, when analyzed together, they can provide high diagnostic value (7, 22). Therefore, in the present study, PCA was performed in an attempt to establish a pattern of cytokines and mediators in blood and pleural fluid capable of discriminating the PlTB and non-TB cases. Our results show that 9 cytokines and biomarkers, measured in blood and pleural fluid, were reduced to two principal components (Table 2). Together, they were able to discriminate the PlTB and non-TB cases and explained 57% of the variation. The cytokines with determinant values were ADA, TGF-β, IP-10, IL-17A, IFN-γ, TNF, IL-4, and IL-2. These results provide new data in the search for new markers capable of differentiating the causes of exudative pleural effusion.

Some limitations should be considered in our study. First, it was conducted in a single-center, imposing a validation in other reference centers and in different populations. Another consideration is regarding the relatively low number of patients included per group. However, patients were included prospectively, in a real routine of clinical practice in a tertiary reference center which reflected in variable clinical characteristics inherent of each group of study, as can be observed in Table 1. Moreover, we have analyzed only exudative cases of pleural effusion, the main confounders in the differential diagnosis of TB. Also, we have excluded transudative cases which could add some bias in our analysis.

In summary, the analysis of a panel of inflammatory mediators previously highlighted in the TB literature was useful to provide new hypotheses and better comprehension about microenvironment of the pleural cavity during the immunopathology of Mtb infection. In addition, the screening in pleural fluid identified biomarkers with high potential for use alone or in combination, which is able to increase the sensitivity of diagnosis and prompt the TB treatment, especially in cases of hard identification and distinction by conventional diagnostic methods.

## Acknowledgments

We would like to thank the physicians of Tuberculosis Outpatient Clinics of HUPE/UERJ, which contributed to the medical care of the patients included in the study.

## Author Contributions

### Conceptualization

Luciana Silva Rodrigues.

### Data curation

Vinicius da Cunha Lisboa, Raquel da Silva Corrêa, Isabelle Ramos Lopes.

### Formal analysis

Vinicius da Cunha Lisboa, Raquel da Silva Corrêa, Marcelo Ribeiro-Alves.

### Funding acquisition

Thaís Porto Amadeu, Rogério Lopes Rufino Alves, Luciana Silva Rodrigues.

### Investigation

Vinicius da Cunha Lisboa, Thiago Thomaz Mafort, Ana Paula Gomes dos Santos, Rogério Lopes Rufino Alves, Luciana Silva Rodrigues.

### Methodology

Vinicius da Cunha Lisboa, Raquel da Silva Corrêa, Isabelle Ramos Lopes.

### Project administration

Luciana Silva Rodrigues.

### Resources

Rogério Lopes Rufino Alves, Luciana Silva Rodrigues.

### Software

Vinicius da Cunha Lisboa, Marcelo Ribeiro-Alves.

### Supervision

Luciana Silva Rodrigues.

### Validation

Marcelo Ribeiro-Alves.

### Writing – original draft

Vinicius da Cunha Lisboa, Raquel da Silva Corrêa, Marcelo Ribeiro-Alves, Luciana Silva Rodrigues.

### Writing – review & editing

Vinicius da Cunha Lisboa, Marcelo Ribeiro-Alves, Luciana Silva Rodrigues.

## Data Availability Statement

All relevant data are within the manuscript and its Supporting Information files.

